# Comparative genomic analyses reveal diverse virulence factors and antimicrobial resistance mechanisms in clinical *Elizabethkingia meningoseptica* strains

**DOI:** 10.1101/668061

**Authors:** Shicheng Chen, Marty Soehnlen, Jochen Blom, Nicolas Terrapon, Bernard Henrissat, Edward D. Walker

**Author notes:** Corresponding author: (SC).

## Abstract

Three human clinical isolates of bacteria (designated strains Em1, Em2 and Em3) had high average nucleotide identity (ANI) to *Elizabethkingia meningoseptica*. Their genome sizes (3.89, 4.04 and 4.04 Mb) were comparable to those of other *Elizabethkingia* species and strains, and exhibited open pan-genome characteristics, with two strains being nearly identical and the third divergent. These strains were susceptible only to trimethoprim/sulfamethoxazole and ciprofloxacin amongst 16 antibiotics in minimum inhibitory tests. The resistome exhibited a high diversity of resistance genes, including 5 different lactamase- and 18 efflux protein-encoding genes. Forty-four genes encoding virulence factors were conserved among the strains. Sialic acid transporters and curli synthesis genes were well conserved in *E. meningoseptica* but absent in *E. anophelis* and *E. miricola. E. meningoseptica* carried several genes contributing to biofilm formation. 58 glycoside hydrolases (GH) and 25 putative polysaccharide utilization loci (PULs) were found. The strains carried numerous genes encoding two-component system proteins (56), transcription factor proteins (187~191), and DNA-binding proteins (6~7). Several prophages and CRISPR/Cas elements were uniquely present in the genomes.

## Introduction

*Elizabethkingia meningoseptica* is a Gram-negative, non-fermenting, aerobic bacterium occurring in soil, water, plants and animals [1]. *E*. *meningoseptica* normally does not cause infection and disease in healthy humans, but it is a serious causative agent of nosocomial pneumonia, neonatal meningitis, bacteremia, and endocarditis in immuno-compromised patient populations [2–4]. Infections are mostly acquired in hospitals as bacteria are often recovered from medical apparatus and reagents including tap water, disinfection fluid, ventilators, hemodialysis equipment and catheters; however, sporadic, community-acquired infections were also reported [3–6]. Direct transmission pathways for *E*. *meningoseptica* remain largely unknown [7].

Clinical manifestations of *E*. *meningoseptica* infections are similar to those of other *Elizabethkingia* species or other bacteria though some variations exist [8, 9]. *Elizabethkingia* infections in neonates or immuno-compromised patients are usually associated with a poor outcome [2, 3]. One of the most challenging issues is that routine morphological, biochemical, and molecular tests cannot accurately identify *Elizabethkingia* species [10]. Sequencing of the 16S rRNA gene often fails to provide sufficient resolution to differentiate various *Elizabethkingia* species which may show different antibiotic resistance properties [10, 11]. For example, recent studies have demonstrated that *E. anophelis* or *E. meningoseptica* were frequently misidentified; it is not surprising that some *E. meningoseptica* infections were actually reported to be caused by *E. anophelis* [12]. Thus, accurate and complementary diagnostic methods need to be developed such as MALDI-ToF or genome sequencing.

*Elizabethkingia* are typically highly resistant to antimicrobials including extended-spectrum beta-lactams, tetracycline, aminoglycosides, and chloramphenicol [13]. Nevertheless, *E. meningoseptica*-infections are empirically treated with antibiotics used for Gram-positive bacteria [14]. A growing number of studies demonstrate that *Elizabethkingia* species (even the same species) isolated from different geographical regions have different antibiotic susceptibilities, showing that there is a complex antimicrobial resistance spectrum in *Elizabethkingia* [3, 13, 15]. Consequently, the high mortality rates (up to 50%) reported for immunosuppressed patients may, at least partially, be the result of a delay in identifying the appropriate antibiotic therapy [3, 10, 13]. Thanks to next generation sequencing technology, the studies on the antimicrobial resistance mechanisms have significantly progressed. Recently, comparative genome analysis in *Elizabethkingia* has been conducted to discover the breadth of antibiotic susceptibility and epidemiological features [16–18]. However, most of the studies are limited to *E. anophelis* and focused on clinical cases, antimicrobial susceptibility patterns, or characterization of epidemic outbreaks [16–18]. Genome analyses and physiological studies have not been conducted in depth for *E*. *meningoseptica*.

The aim of this study was to analyze virulence, antibiotic resistance, environmental survival, and adaption mechanisms through genomic, resistome, and antibiotic susceptibility analyses. With the completion of genome sequencing, assembly, and annotation of *E. meningoseptica* strains, we examined gene repertoire and genetic diversity in comparison with other *Elizabethkingia* species. Further, the regulatory systems, sugar utilization systems, virulence factors and antibiotic resistance genes were analyzed and compared to those in the selected *Elizabethkingia*. Collectively, our study offers the opportunity to explore their virulence, antibiotic resistance, environmental survival, and adaptation mechanisms.

## Materials and Methods

### Culture conditions

*E. meningoseptica* strains, designated here Em1, Em2 and Em3, were isolated from patients in Michigan (Table 1). *E. meningoseptica* strains were grown aerobically in tryptic soy broth (TSB) broth at 30°C. Bacto agar (Difco, Detroit, MI) was added to a final concentration of 20 g/liter. The sheep blood agar (SBA) was purchased from Thermo Scientific (Waltham, MA).

**Table 1.** General features of selected *Elizabethkingia* genomes.

| Strain | Original<br>region<sup>*a</sup> | Specimen<sup>*b</sup> | Size<br>(Mb) | GC<br>% | Protein | Gene | Pseudogene |
| --- | --- | --- | --- | --- | --- | --- | --- |
| ***E. meningoseptica*** |  |  |  |  |  |  |  |
| EM1 | USA | Tracheal<br>exudate | 4.04 | 36.4 | 3,557 | 3,656 | 44 |
| EM2 | USA | Whole blood | 3.89 | 36.4 | 3,403 | 3,483 | 27 |
| EM3 | USA | Sputum | 4.04 | 36.4 | 3,556 | 3,656 | 45 |
| G4120 | France | Urine | 4.00 | 36.4 | 3,519 | 3,630 | 47 |
| NBRC 12535 | USA | Spinal fluid | 3.84 | 36.4 | 3,396 | 3,459 | 20 |
| 61421 PRCM | China | NA | 3.96 | 36.4 | 3,466 | 3,546 | 31 |
| G4076 | UK | NA | 3.87 | 36.5 | 3,406 | 3,483 | 14 |
| CSID_3000516977 | USA | NA | 3.91 | 36.4 | 3,421 | 3,492 | 26 |
| 58-80 | Israel | Tracheal<br>exudate | 3.84 | 36.6 | 3,359 | 3,484 | 76 |
| ***E. anophelis*** |  |  |  |  |  |  |  |
| CSID_3015183678 | USA | NA | 3.93 | 35.8 | 3,473 | 3,578 | 35 |
| NUHP1 | Singapore | NA | 4.37 | 35.6 | 3,912 | 4,039 | 58 |
| Ag1 | USA | Mosquito | 4.09 | 35.5 | 3676 | 3780 | 34 |
| ***E. miricola*** |  |  |  |  |  |  |  |
| BM10 | Korea | Termite | 4.24 | 35.7 | 3,728 | 3,728 | 89 |
| CSID_3000517120 | USA | NA | 4.43 | 35.9 | 3,932 | 4,028 | 45 |

<sup>\*a, b</sup> The information about specimen and sources used for these selected isolates was obtained from BioSample (https://www.ncbi.nlm.nih.gov/genome/genomes/14625). NA, not available.

### Antimicrobial susceptibility testing (AST)

Cultures were freshly grown on SBA overnight at 37°C and cell suspensions adjusted in saline solution (0.9%) to a turbidity equivalent of 0.5 McFarland standard. The VITEK 2 microbial ID/AST testing system (version 07.01) with the GNP 70 antibiotic susceptibility cards (bioMerieux) was used to determine the minimum inhibitory concentration (MIC) and classification into resistance phenotypes. MIC results were interpreted according to Clinical and Laboratory Standards Institute criteria (CLSI) [19].

### Genomic DNA preparation, sequencing and assembly

DNA was isolated using a Wizard Genomic DNA Purification Kit (Promega, Madison). The concentration of genomic DNA was measured using a Nanodrop2000 UV-Vis Spectrophotometer (Thermo scientific) and Qubit DNA assay kit. DNA integrity was evaluated by agarose gel assay (1.5%, w/v).

NGS libraries were prepared using the Illumina TruSeq Nano DNA Library Preparation Kit. Completed libraries were evaluated using a combination of Qubit dsDNA HS, Caliper LabChipGX HS DNA and Kapa Illumina Library Quantification qPCR assays. Libraries were combined in a single pool for multiplex sequencing and the pool was loaded on one standard MiSeq flow cell (v2) and sequencing performed in a 2×250 bp, paired end format using a v2, 500 cycle reagent cartridge. Base calling was done by Illumina Real Time Analysis (RTA) v1.18.54 and output of RTA was demultiplexed and converted to FastQ format with Illumina Bcl2fastq v1.8.4. The genomes were assembled into contiguous sequences using SPAdes version 3.9 following the manual as described previously[20], then short and low-coverage contigs were filtered out.

### Genome annotation

Annotation of the assembled genome sequences for Em1, Em2 and Em3 was submitted to the NCBI Prokaryotic Genome Automatic Annotation Pipeline (PGAAP). The predicted CDSs were translated and analyzed against the NCBI non-redundant database, Pfam, TIGRfam, InterPro, KEGG and COG. Additional gene prediction and manual revision was performed by using the Integrated Microbial Genomes and Microbiome Samples Expert Review (IMG/MER) platform.

### Bioinformatics

Functional categorization and classification for predicted ORFs were performed by RAST server-based SEED viewer [21]. Multi-drug resistance genes were predicted in the Comprehensive Antibiotic Resistance Database [22]. Prophage prediction was done with PHAST [23] and Clustered Regularly Interspaced Short Palindromic Repeats (CRISPR) were predicted by CRISPRfinder [24]. For genome similarity assessment, average nucleotide identity (ANI) was computed using ANI calculator (https://www.ezbiocloud.net/tools/ani).

The pan genome, core genome, singletons and specific genes of Em1, Em2 and Em3 were characterized by comparing the representative *Elizabethkingia* genomes using EDGAR 2.0 [25]. The development of pan genome and core genome sizes was approximated using the methods proposed by Tettelin et al [26]. The core genome calculated by EDGAR 2.0 was the reference used to infer a phylogeny. The 27,144 amino acid sequences (2,088 per genome) of the core genome were aligned set-wise using MUSCLE v3.8.31 [27], resulting in a large multiple alignment with 9,015,188 amino acid residues in total (693,476 per genome). This large alignment was used to construct a phylogenetic tree using the neighbor-joining method as implemented in the PHYLIP package [28].

The regulatory elements were predicted using p2rp with default settings (http://www.p2rp.org). Bacterial protein localization prediction was conducted with tool PSORTb version 3.0.2 (http://www.psort.org/psortb/). Carbohydrate active enzyme families, including enzymes of glycan assembly (glycosyltransferases, GT) and deconstruction (glycoside hydrolases, GH, polysaccharide lyases, PL, carbohydrate esterases, CE), were semi-manually annotated using the Carbohydrate Active Enzyme (CAZy) database curation pipelines [29]. Polysaccharide-utilization loci (PUL) was predicted as described in the PULDB database (www.cazy.org/PULDB/).

### Biofilm formation

Biofilm formation was evaluated using a standard assay with crystal violet staining as previously described, [30] with modifications as follows. Em1, Em2 and Em3 were grown in TSB broth to the mid-exponential phase. The cultures were diluted in TSB broth, and 100 µL were deposited in wells of 96-well microtiter polystyrene plates with flat bottoms. The negative controls were TSB medium without inoculum. The plate was incubated at 28°C under static condition for 24 h. The supernatants were discarded, the wells were washed twice with 200 µL of sterile distilled water and then 150 µL of 1% (w/v) crystal violet was added to each well. After 30 min, excess stain was removed by washing the wells four times with 200 µL of sterile distilled water and the stain bound to adherent cells was subsequently released by adding 100 µL of absolute ethanol. The biofilm formation was determined by measuring the OD595 nm using the plate reader.

### Accession of the genome sequences

The data from these Whole Genome Shotgun projects have been deposited at DDBJ/ENA/GenBank under accession numbers MCJH00000000, MDTZ00000000 and MDTY00000000 for Em1, Em2 and Em3, respectively. The BioProject designations for this project are PRJNA336273, PRJNA339645 and PRJNA338129, and BioSample accession numbers are SAMN05507161, SAMN05601445 and SAMN05521514 for Em1, Em2 and Em3, respectively.

### Statistical analyses

Statistical analyses were performed using SAS (version 9.2; SAS Institute, Cary, NC).

## Results

### Genome features and phylogenetic inferences

The assembly of strain Em1, Em2 and Em3 contained 10, 14 and 11 contigs with a size of 4.04, 3.89 and 4.04 Mbp, respectively (Table 1), which is congruent with the genome size (ranging from 3.84 to 4.04 Mbp) among the selected *E. meningoseptica* strains. The Em1, Em2 and Em3 genomes included 3,656, 3,483 and 3,656 coding sequences (CDS) and 55, 53 and 55 RNA genes, respectively. The GC content in the three *E. meningoseptica* strains (Table 1) was ca. 36.4%, consistent with other *E. meningoseptica* isolates (36.2~36.6%); collectively, the average GC content in *E. meningoseptica* genomes (36.4%, n=26) was slightly higher than that in *E. anophelis* (average 35.6%, n=64) or *E. miricola* (35.9%, n=17) (S1 Fig). No plasmid(s) sequences were found. 264 subsystems were identified by RAST analysis (S2 Fig).

ANI values indicated that the three isolates belonged to *E. meningoseptica* species as they were more than 99% identical to the type strain *E. meningoseptica* ATCC 13253 (Table 2). However, ANI values were low (<80%) in comparisons with *E. anophelis* or *E. miricola* (Table 2). The difference in overall genome features between Em1 and Em3 was negligible (Table 1). Phylogenetic trees (S3 Fig) showed that Em1 and Em3 were clustered more closely to *E. meningoseptica* CSID 30005163359 and G58-80. However, the phylogenetic placement of Em2 was closer to *E. meningoseptica* CSID 3000516465, which formed a different clade from the other two strains (S3 Fig).

**Table 2.** Average nucleotide identity dendrogram for the selected *Elizabethkingia* spp*.

|  |  | Em2 | 61421 PRCM | G4120 | 58-80 | Em1 | Em3 | CSID_3000515919 | CSID_3000516977 | KC1913 | Ag1 | NUHP1 | CSID_3015183678 | BM10 | CSID_3000517120 |
| --- | --- | --- | --- | --- | --- | --- | --- | --- | --- | --- | --- | --- | --- | --- | --- |
| *E. meningoseptica* | Em2 | 100 |  |  |  |  |  |  |  |  |  |  |  |  |  |
|  | 61421 PRCM | 99.76 | 100 |  |  |  |  |  |  |  |  |  |  |  |  |
|  | G4120 | 99.75 | 99.89 | 100 |  |  |  |  |  |  |  |  |  |  |  |
|  | 58-80 | 99.37 | 99.31 | 99.35 | 100 |  |  |  |  |  |  |  |  |  |  |
|  | Em1 | 99.18 | 99.17 | 99.32 | 99.36 | 100 |  |  |  |  |  |  |  |  |  |
|  | Em3 | 99.17 | 99.16 | 99.3 | 99.36 | 99.98 | 100 |  |  |  |  |  |  |  |  |
|  | CSID_3000515919 | 98.54 | 98.38 | 98.6 | 98.67 | 98.52 | 98.51 | 100 |  |  |  |  |  |  |  |
|  | CSID_3000516977 | 98.22 | 98.21 | 98.29 | 98.36 | 98.28 | 98.24 | 98.25 | 100 |  |  |  |  |  |  |
|  | KC1913 | 98.27 | 98.3 | 98.3 | 98.39 | 98.25 | 98.24 | 98.3 | 99.09 | 100 |  |  |  |  |  |
| *E. anophelis* | Ag1 | 79.67 | 79.71 | 79.7 | 79.91 | 79.66 | 79.67 | 79.97 | 79.81 | 79.82 | 100 |  |  |  |  |
|  | NUHP1 | 79.76 | 80.08 | 79.74 | 79.77 | 79.69 | 79.77 | 79.9 | 79.74 | 79.78 | 97.83 | 100 |  |  |  |
|  | CSID_3015183678 | 79.76 | 79.86 | 79.81 | 79.92 | 79.7 | 79.82 | 79.78 | 79.79 | 79.86 | 98.19 | 97.99 | 100 |  |  |
| *E. miricola* | BM10 | 79.84 | 79.83 | 79.77 | 80.01 | 79.78 | 79.8 | 79.97 | 79.78 | 79.78 | 90.93 | 90.41 | 90.93 | 100 |  |
|  | CSID_3000517120 | 79.67 | 79.97 | 79.63 | 80.29 | 79.6 | 79.58 | 79.66 | 79.67 | 79.74 | 91.13 | 90.73 | 91.27 | 92.37 | 100 |

\*ANI matrix was generated using ANI calculator (https://www.ezbiocloud.net/tools/ani).

### Gene repertoire of *E. meningoseptica*

The core genome and pan-genome were sorted and used for gene repertoire analysis in selected *E. meningoseptica* genomes (Fig 1A, 1B). Core genome analysis showed that the number of shared genes decreased with addition of the input genomes (see Fig 1B). Overall, *E. meningoseptica* displayed an open pan-genome feature because new genes appeared when more sequenced genomes were added to the analysis (Fig 1A). *E. meningoseptica* Em1 and Em3 shared at least 3556 genes; only 3 and 5 genes were uniquely present in strains Em1 and Em3, respectively, supporting that Em1 and Em2 are very similar (Fig S4). Isolate Em2 shared 3280 and 3278 genes with Em1 and Em3, respectively, while there were 125 and 127 unique genes in Em2 (S4 Fig). Moreover, *E. meningoseptica* Em2 shared 3,184, 3,326, 3,106, 3,322, 3,173 and 3,788 common genes with strains G4076, G4120, NBRC 12535, and 61421 PRCM, respectively (Fig 2A). These genes shared in common accounted for approximately 93.5%, 97.7%, 91.2%, and 97.5% of the encoding genes of Em2, respectively; taken together, Em2 and these four strains shared 3,076 genes (Fig 2A). However, *E. meningoseptica* Em2 shared far fewer genes with *E. anophelis* strains (2,772 to 2,823) and *E. miricola* (2,789 to 2,890), accounting for less than 86% of the total encoding genes (Fig 2B).

**Fig 1.**
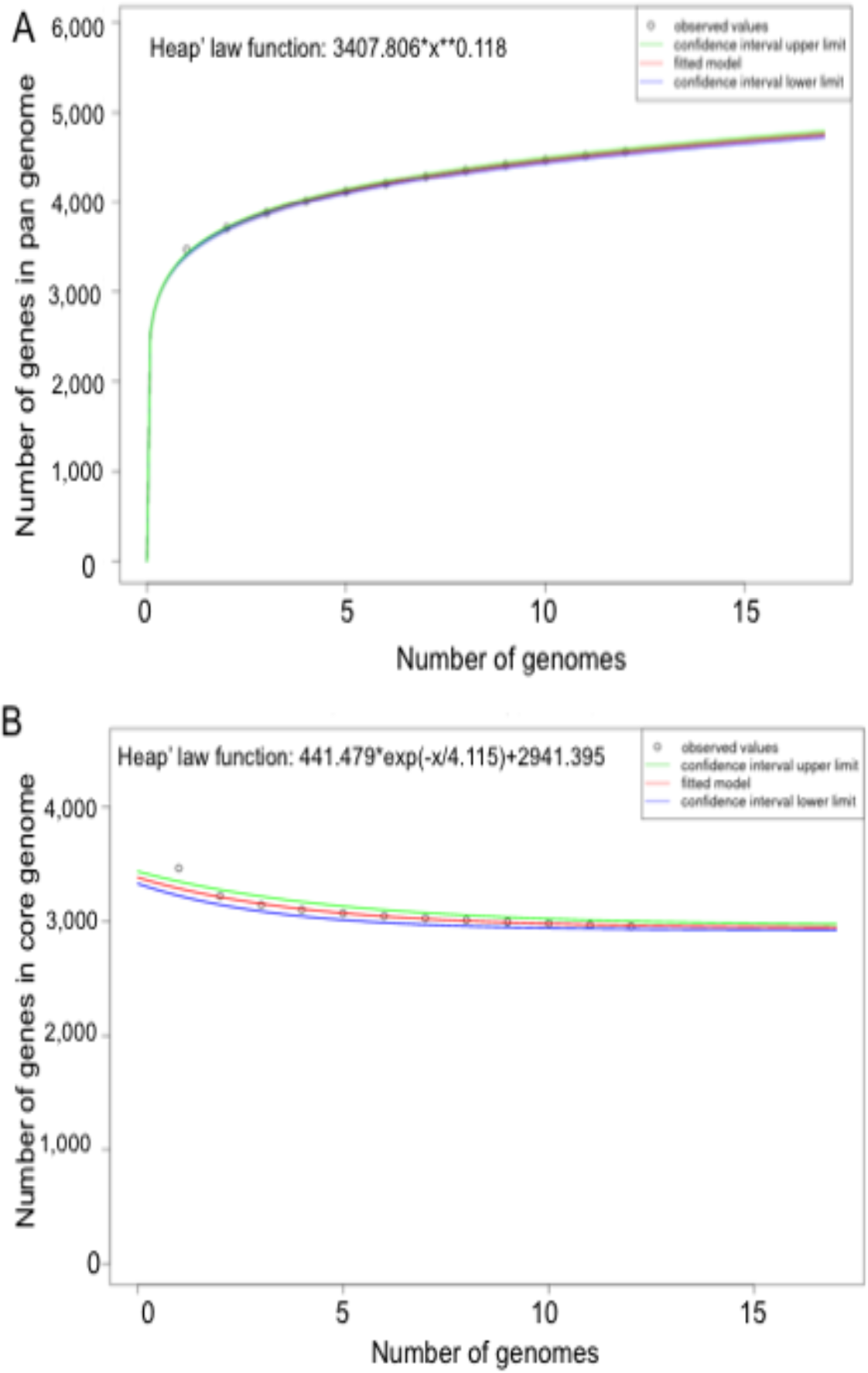
Pan, core, and singleton genome evolution according to the number of selected *Elizabethkingia* genomes. (A) Number of genes (pan-genome) for a given number of genomes sequentially added. The pan development plot was generated for the following genomes: *E. meningoseptica* EM2 (NZ_MDTZ01000014), *E. meningoseptica* NV2016 (NZ_FRFB01000021), *E. meningoseptica* G4120 (NZ_CP016378), *E. meningoseptica* CSID3000516359 (NZ_MAHC01000017), *E. meningoseptica* CCUG214 (NZ_FLSV01000010), *E. meningoseptica* CSID_3000515919 (NZ_MAGZ01000024), *E. meningoseptica* CSID_3000516535 (NZ_MAHF01000020), *E. meningoseptica* ATCC13253 (NBRC_12535), *E. meningoseptica* EM3 (NZ_MDTY01000011), *E. meningoseptica* CIP111048 (NZ_FTPF01000022), *E. meningoseptica* 61421PRCM (NZ_MPOG01000010), *E. meningoseptica* EM1 (NZ_MCJH01000010), *E. meningoseptica* 58_80 (NZ_FTRA01000043). (B) Number of shared genes (core genome) as a function of the number of genomes sequentially added. The genomes used for generating the core genome development plot were the same as listed in (A).

**Fig 2.**
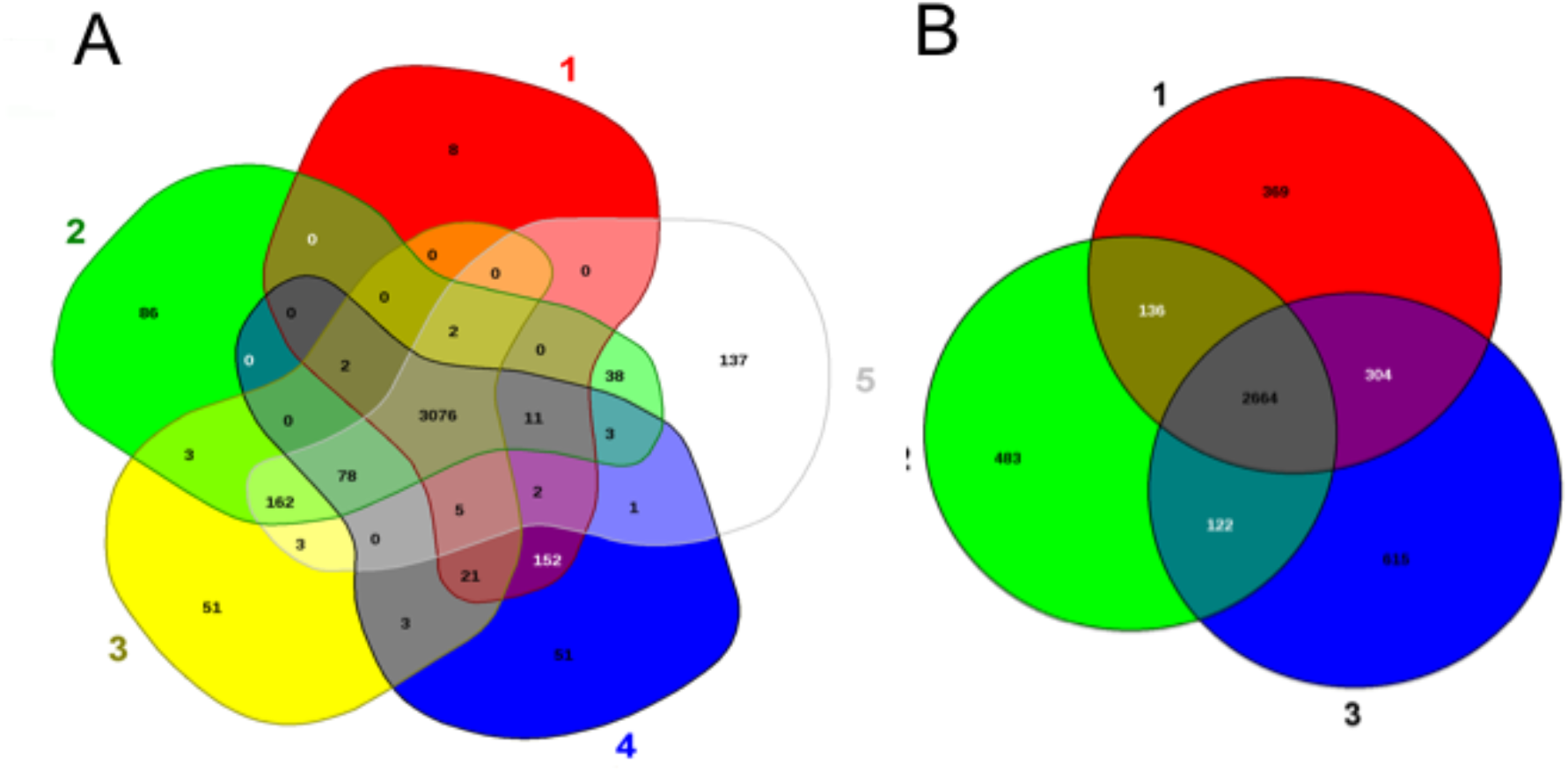
Venn diagram of shared and unique genes in selected *Elizabethkingia*. The unique and shared genome among the selected strains was determined by a dual cutoff of 30% or greater amino acid identity and sequence length coverage of more than 70%. EDGAR was used for Venn diagrams. A) 1: *E. anophelis* CSID_3015183678, 2: *E. meningoseptica* Em2 and 3: *E. miricola* BM10. B) 1: *E. meningoseptica* ATCC 13253, 2: *E. meningoseptica* 61421 PRCM, 3: *E. meningoseptica* Em2, *E. meningoseptica* G4076, and *E. meningoseptica* G4120.

### Antimicrobial susceptibility and antibiotic resistance gene analysis

Susceptibilities of the *E. meningoseptica* isolates and ATCC13253 with corresponding MICs to tested antibiotics showed very similar antibiotic susceptibility spectra with minor discrepancies across strains (Table 3). The strains were highly resistant to 13 of 16 antimicrobial reagents, showing that they were multi-drug resistance strains (Table 3).

**Table 3.** Antibiotic susceptibility test (AST) in the selected *E. menigoseptica*.

| Antibiotic class | Selected antibiotics | *E. meningoseptica* |  |  |  |
| --- | --- | --- | --- | --- | --- |
|  |  | Em1 | Em2 | Em3 | ATCC13253 |
| Aminoglycosides | Amikacin | ≥64 (R) | ≥64 (R) | ≥64 (R) | ≥64 (R) |
|  | Gentamicin | ≥16 (R) | ≥16 (R) | ≥16 (R) | ≥16 (R) |
|  | Tobramycin | ≥16 (R) | ≥16 (R) | ≥16 (R) | ≥16 (R) |
| β-lactams | Aztreonam | ≥64 (R) | ≥64 (R) | ≥64 (R) | ≥64 (R) |
|  | Ampicillin | ≥32 (R) | ≥32 (R) | ≥32 (R) | ≥32 (R) |
|  | Ampicillin/Sulbactam | ≥32 (R) | ≥32 (R) | ≥32 (R) | ≥32 (R) |
|  | Piperacillin | ≥64 (R) | ≥64 (R) | ≥64 (R) | ≥64 (R) |
|  | Ceftriaxone | ≥64 (R) | ≥64 (R) | ≥64 (R) | ≥64 (R) |
|  | Piperacillin/Tazobactam | ≥128 (R) | ≥128 (R) | ≥128 (R) | ≥128 (R) |
|  | Cefazolin | ≥64 (R) | ≥64 (R) | ≥64 (R) | ≥64 (R) |
|  | Cefotaxime | ≥32 (R) | ≥32 (R) | ≥32 (R) | ≥32 (R) |
|  | Meropenem | ≥16 (R) | ≥8 (I) | ≥16 (R) | ≥16 (R) |
| Sulfonamides | Trimethoprim/Sulfamethoxazole | 40 (S) | 40 (S) | 40 (S) | 40 (S) |
| Quinolone | Ciprofloxacin | 1 (S) | 1 (S) | 1 (S) | 1 (S) |
| Glycylcycline | Tigecycline | 4 (I) | 8 (R) | 4 (I) | 8 (R) |
| Others | Nitrofurantoin | 128(R) | 512 (R) | 256 (R) | 256 (R) |

The values were minimum inhibitory concentration (µg/ml) determined by the VITEK as described in Methods and Materials; S, I or R in the parenthesis stands for sensitive (S), intermediately sensitive (I), resistant (R). The sensitive and intermediately sensitive susceptibility were highlighted with red font.

Twenty-two genes encoding enzymes/proteins conferring resistance to antimicrobial reagents were found by CARD and RAST SEED subsystem (Table 4). Up to 5 lactamase genes encoding β-lactamases (EC 3.5.2.6), metal-dependent hydrolases (superfamily I), class C β-lactamases/penicillin binding protein and other β-lactamases were predicted in Em2, which possibly contributed to the intrinsic resistance to six β-lactam drugs tested in this study (Table 3). One fluoroquinolone resistance gene (*gyrB*) and three genes (*otr*A, *tetO* and *tetBP*) by RAST analysis possibly involved in tetracycline resistance were discovered. Moreover, 18 multidrug resistance efflux pumps (Table 4) were revealed, which may confer non-specific resistance (Table 3).

**Table 4.**
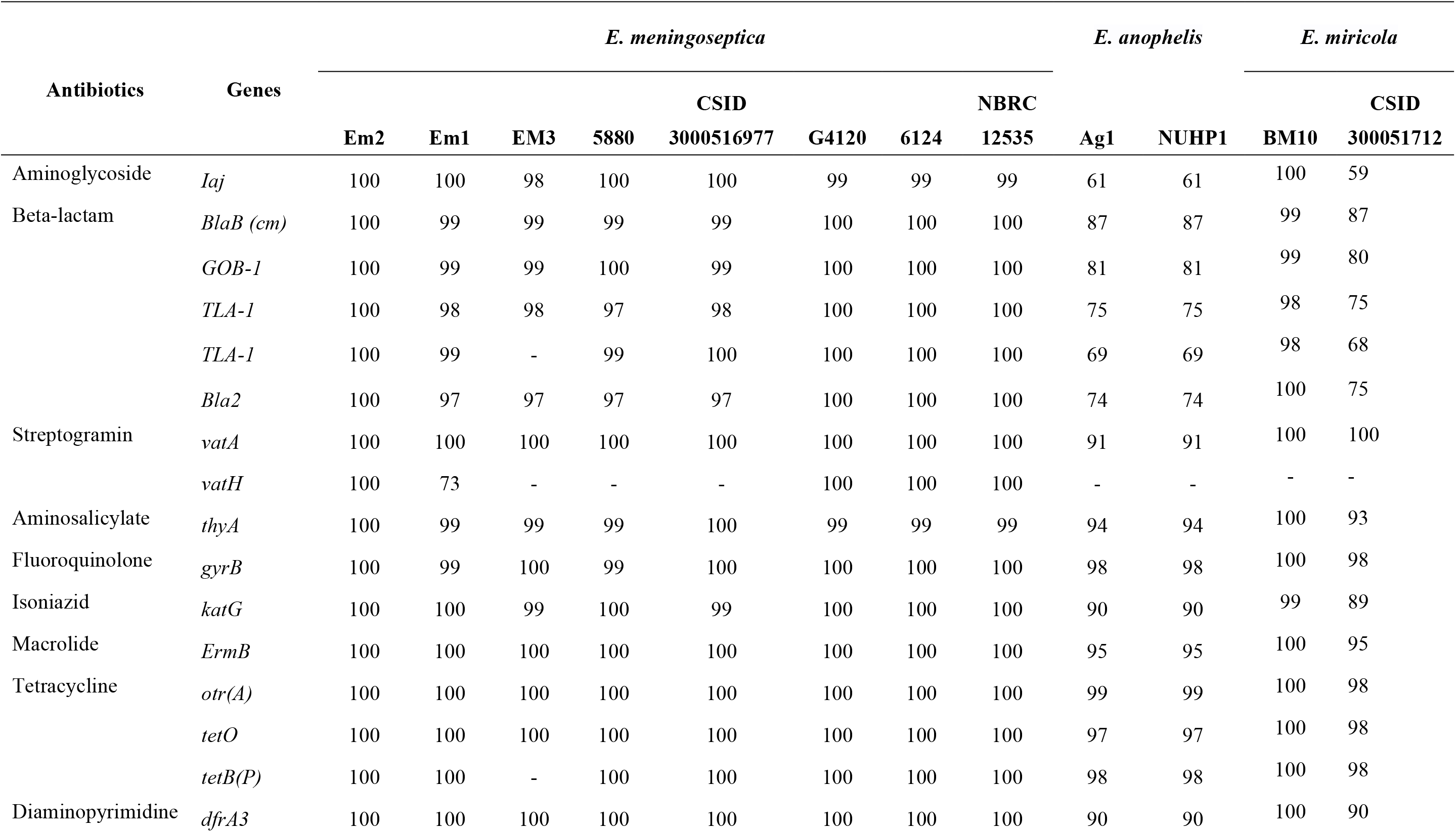
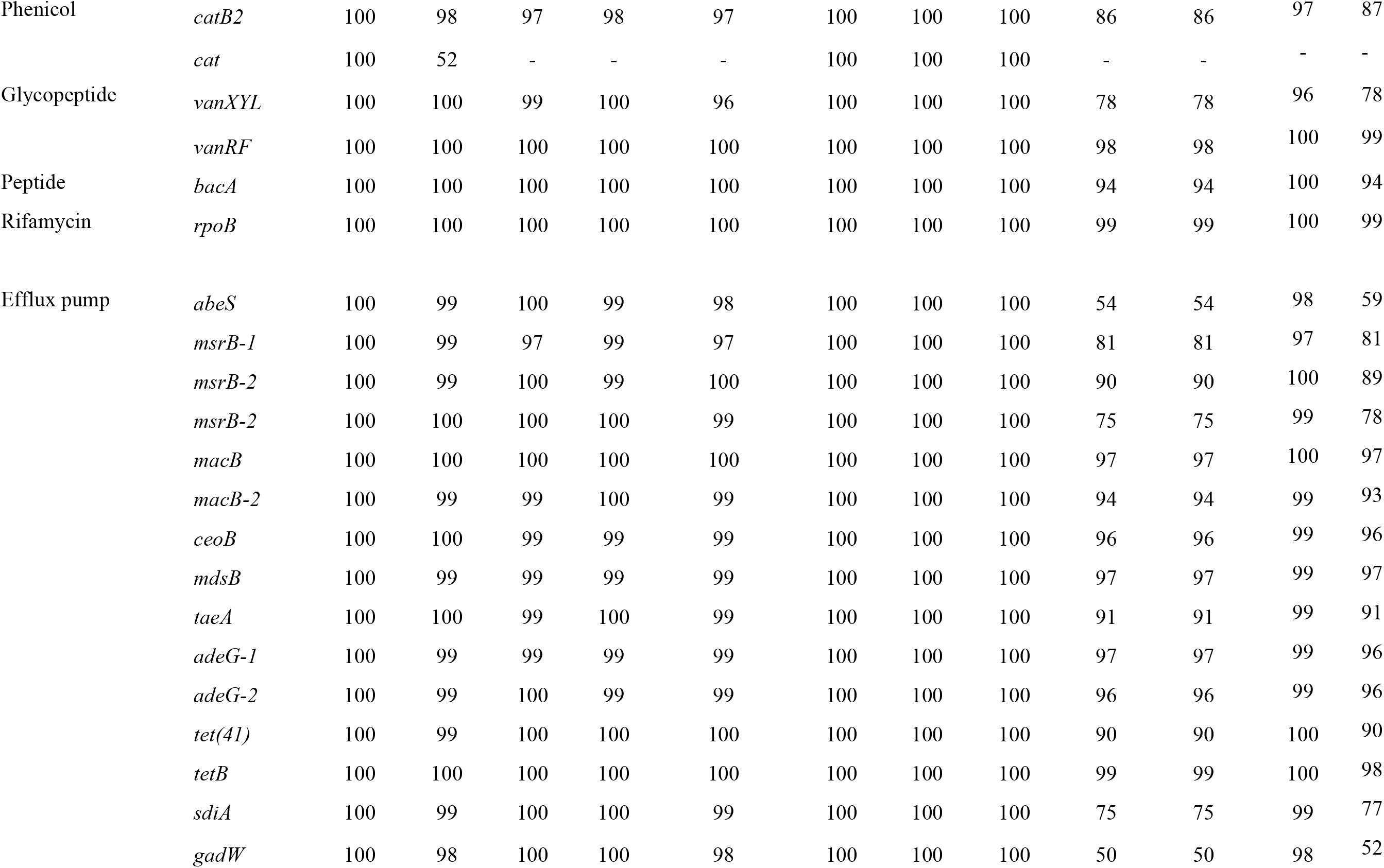
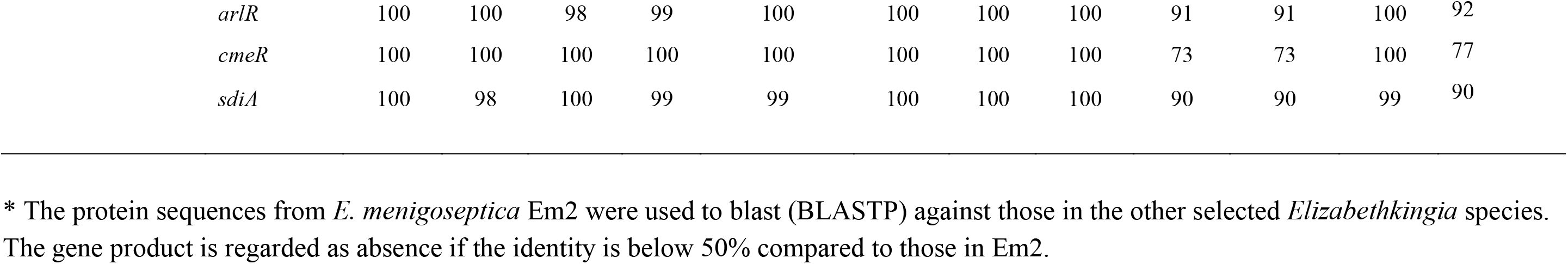
Comparison of similarity (% nucleotide sequence) of antimicrobial resistance genes in various *Elizabethkingia*\*.

### Virulence factors predicted in *E. meningoseptica* in comparison to other *Elizabethkingia* spp

The pathogenesis mechanisms in *Elizabethkingia* species remain largely unknown. When PathogenFinder was used to predict the probability of acting as a pathogen for strains Em1, Em2 and Em3, *E. meningoseptica* showed a “negative” result according to the calculated probability values (~0.162) [31]. Instead, *Elizabethkingia* genomes only matched 5 non-pathogenic families (data not shown), which was possibly due to the lack of similar flavobacterial genomes in the PathogenFinder database. However, 766 putative virulence factors in Em2 were successfully predicted by VFDB (cutoff value set to E^−10^, data not shown). Forty-four of them in the bacterial virulence database also showed high identity to the selected *Elizabethkingia*, although there were some minor variations (S1Table). Many virulence genes were predicted to be involved in CME-1 formation, capsule polysaccharide synthesis, lipooligosaccharide (LOS) synthesis, “attacking” enzymes (proteases), superoxide dismutase, catalases, peroxidase, heat shock protein, two-component regulatory system and many others (see S1 Table).

*E. meningoseptica* EM2 carried at least 6 genes involved in sialic acid metabolism predicted by RAST while Em1 and Em2 only possessed 4 of them (Table 5). Three genes encoded glucosamine-6-phosphate deaminase (*nagB*), glucosamine-fructose-6-phosphate aminotransferase (*glmS*) and phosphoglucosamine mutase (*glmB*), which were involved in the *de novo* synthesis pathway of sialic acid. Gene products for sialic acid synthesis were highly conserved (identity > 93%) among the three selected *Elizabethkingia* spp (Table 5). Two copies of sialic acid transporter genes (*neuC*1 *and neuC*2, encoding UDP-N-acetylglucosamine 2-epimerase) were found in strains Em2, G4120 and 61421 PRCM; CSID_3000515919 only carried only one copy (*neuC2*). An *nanH* gene, encoding a candidate sialidase, was predicted as the virulence factor (Table 5). Among the selected *Elizabethkingia*, only *E. meningoseptica* carries *nanH* (Table 5). Moreover, neither *E. miricola* (except strain CSID_3000517120) nor *E. anophelis* had genes encoding sialidase A (*nanH*) or sialic acid transporter genes (*neu* C1 and C2).

**Table 5.**
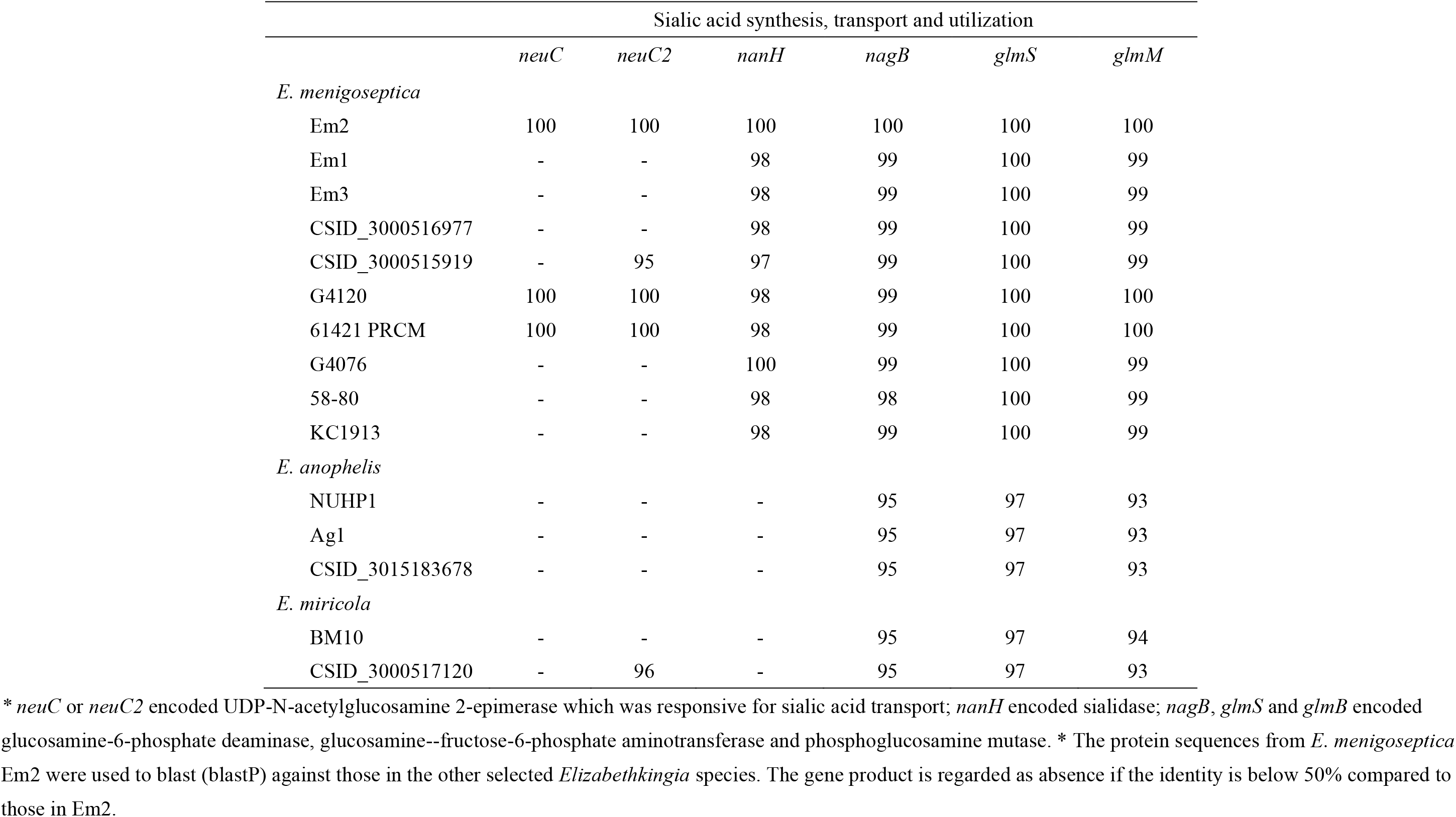
Comparison of selected sialic acid synthesis, transport and utilization genes among *Elizabethkingia*\*.

|  | Sialic acid synthesis, transport and utilization |  |  |  |  |  |
| --- | --- | --- | --- | --- | --- | --- |
|  | *neuC* | *neuC2* | *nanH* | *nagB* | *glmS* | *glmM* |
| *E. meningoseptica* |  |  |  |  |  |  |
| Em2 | 100 | 100 | 100 | 100 | 100 | 100 |
| Em1 | - | - | 98 | 99 | 100 | 99 |
| Em3 | - | - | 98 | 99 | 100 | 99 |
| CSID_3000516977 | - | - | 98 | 99 | 100 | 99 |
| CSID_3000515919 | - | 95 | 97 | 99 | 100 | 99 |
| G4120 | 100 | 100 | 98 | 99 | 100 | 100 |
| 61421 PRCM | 100 | 100 | 98 | 99 | 100 | 100 |
| G4076 | - | - | 100 | 99 | 100 | 99 |
| 58-80 | - | - | 98 | 98 | 100 | 99 |
| KC1913 | - | - | 98 | 99 | 100 | 99 |
| *E. anophelis* |  |  |  |  |  |  |
| NUHP1 | - | - | - | 95 | 97 | 93 |
| Ag1 | - | - | - | 95 | 97 | 93 |
| CSID_3015183678 | - | - | - | 95 | 97 | 93 |
| *E. miricola* |  |  |  |  |  |  |
| BM10 | - | - | - | 95 | 97 | 94 |
| CSID_3000517120 | - | 96 | - | 95 | 97 | 93 |

\* *neuC* or *neuC2* encoded UDP-N-acetylglucosamine 2-epimerase which was responsive for sialic acid transport; *nanH* encoded sialidase; *nagB*, *glmS* and *glmB* encoded glucosamine-6-phosphate deaminase, glucosamine--fructose-6-phosphate aminotransferase and phosphoglucosamine mutase. \* The protein sequences from *E. meningoseptica* Em2 were used to blast (blastP) against those in the other selected *Elizabethkingia* species. The gene product is regarded as absence if the identity is below 50% compared to those in Em2.

The selected *E. meningoseptica* species produced biofilm on plastic surfaces (S5 Fig). Compared to those in strain Em2, gene clusters for capsular polysaccharide synthesis consisting of *cap8E*, *cap8G*, *cap8O*, *cap8D*, *cap4F*, *cap8F*, *cps4D*, *Cj1137c*, *capD* and *cap4D* were only conserved in strains G4120, 61421 PRCM as well as *E. miricola* CSID 3000517120 (S2 Table). Further, the gene products responsive for curli biosynthesis and assembly were highly conserved in *E. meningoseptica* species while they were absent in other *Elizabethkingia* species (S2 Table). On the other hand, the flagellin biosynthesis protein FlgD and flagellar motor protein MotB were found in all of *Elizabethkingia* species (S2 Table). However, *Elizabethkingia* may not produce functional flagella because they lack of most of the flagellin structure proteins (data not shown), which is congruent with the fact that bacteria *Elizabethkingia* are non-motile.

### Polysaccharide utilization loci and carbohydrate active enzymes

*E. meningoseptica* carried 107 (Em2) or 109 (Em1, Em3) CAZyme-encoding genes occupying ca. 3% of the bacterial genome (S3 Table). The predicted CAZyme repertoires in the three *E. meningoseptica* genomes were similar, with 58 glycoside hydrolases (GHs) and two polysaccharide lyases (PLs) distributed across the same families. The main distinction between Em2 and Em1/Em3 was absence of GT2 and GT32 sequences (S3 Table). The CAZyme repertoires of Em1, Em2 and Em3 strains were similar to other *E. meningoseptica* strains and showed the same features which distinguished *E. meningoseptica* from other *Elizabethkingia* species (S3 Table). Notably, *E. meningoseptica* strains did not encode single copies of GH1 (β-glycosidase), GH5 (subfamily 46) and CBM6 (b-glucan binding) genes systematically found in other species, and had a reduced number of genes encoding enzymes from families GH95 and GH130 (S3 Table). However, *E. meningoseptica* strains specifically encoded a GH33 (sialidase) and an extra GH13 protein. Analysis revealed that *E. meningoseptica* utilized a battery of carbon sources, including D-maltose, D-trehalose, D-gentibiose, D-melibiose, D-glucose, D-mammose, D-fructose, D-fucose, D-mannitol and D-glycerol[11]. Strains Em1, Em2 and Em3 contained 25 loci encoding SusC-SusD homologs in their respective genomes [32], with only four PULs containing GHs. These numbers were similar to those found in other *E. meningoseptica* (23-24 SusCD loci; 5 PULs with GHs) and *E. anophelis* (25 SusCD loci and 4-5 PULs with GHs per genome; n=17) strains. The three clinical strains did not exhibit the PUL that contains the gene encoding the GH13_5 enzyme (α-amylase; conserved but translocated elsewhere in the genome), that was conserved in all *Elizabethkingia* genomes analyzed, including other *E. meningoseptica* strains. However, all six *E. meningoseptica* strains exhibited a specific PUL encoding a GH30_3 enzyme (β-1,6-glucanase) not present in other *Elizabethkingia* genomes (S4 Table). They also harbor an additional GH13 gene in the *Elizabethkingia*-conserved PUL with GH63-GH97 which likely targets storage polysaccharides like glycogen and starch, as in the prototypic starch utilization system [33], and an additional GH16 gene in the *Elizabethkingia*-conserved PUL that encodes a GH29 appended to a CBM32 (S4 Table). Interestingly, the only GH-containing PUL conserved in all *Elizabethkingia* genomes was also widely distributed across the Bacteroidetes phylum, and encoded enzymes from families GH3 (various b-glycosidases) and GH144 (β-1,2-glucanase) which suggests action on bacterial β-1,2-glucans.

### Regulatory systems in *Elizabethkingia* spp

The Em1 genome encoded 56 two-component system proteins, 191 transcription factor proteins, and 7 other DNA-binding proteins (Table 6). Em3 had the same regulatory systems as those in Em1 (Table 6). The Em2 genome encoded 56 predicted two-component system proteins, 187 transcription factor proteins, and 6 other DNA-binding proteins. It seemed that strain Em1 or Em3 had more regulatory capacity than strain Em2 primarily due to the number of transcriptional regulators (TRs), one-component systems (OCS), and DNA-binding proteins (ODP) proteins. Most *E. meningoseptica* strains possessed similar numbers of total regulatory protein (ranging from 246 to 254) and components (Table 6). Comparable regulatory systems occurred in *E. anophelis*, although there were more ODPs in *E. anophelis* CSID_3015183678 and Ag1. Remarkably, the total number of TRs (>140) in *E. miricola* was much higher than those in *E. meningoseptica* and *E. anophelis* (Table 6). Particularly, the TRs mostly accounted for the higher regulatory systems due to increasing AraC family transcription factors (Table 6). AraC family regulators control a variety of cellular processes including carbon metabolism, stress responses and virulence. Moreover, the two component systems and DNA-binding proteins were more abundant in the three selected *E. miricola* strains than those in *E. meningoseptica*.

**Table 6.**
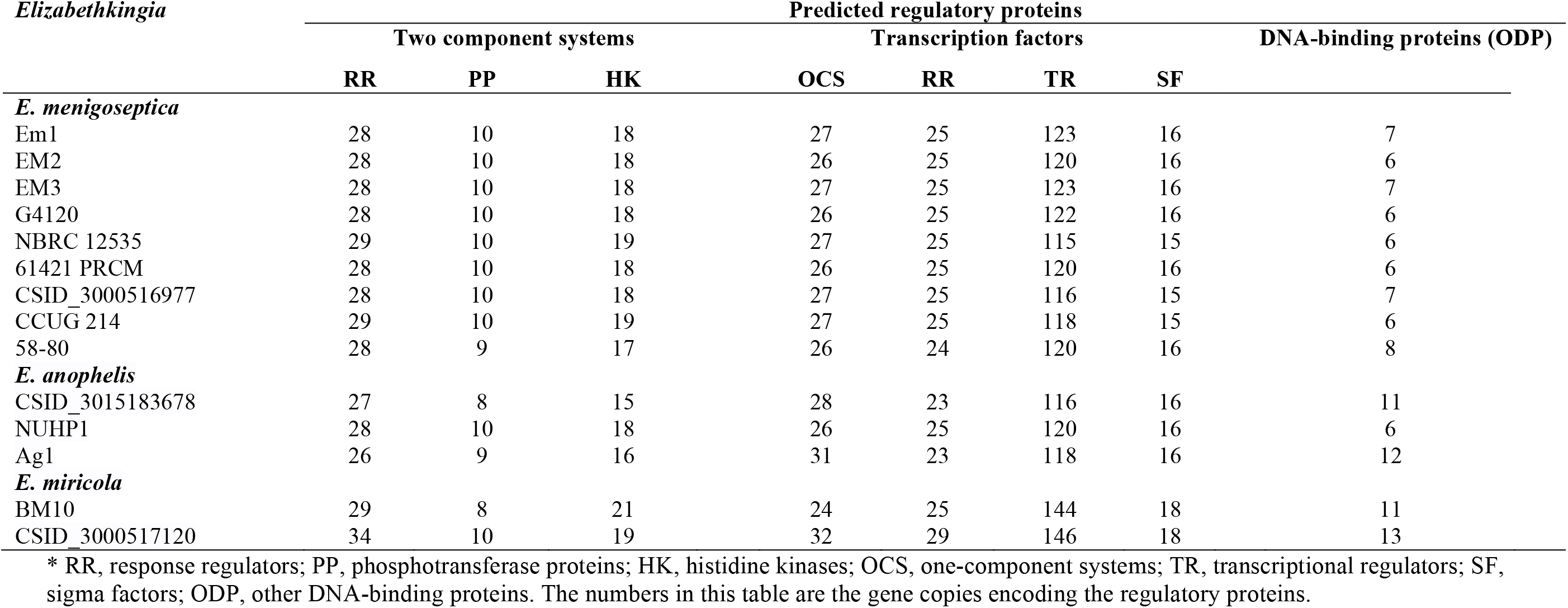
Predicted regulatory systems in *Elizabethkingia*\*.

### CRISPR-Cas systems and prophages

The clustered regularly interspaced short palindromic repeat (CRISPR)/CRISPR-associated protein (CRISPR/Cas) system was predicted in the Em1 and Em3 genome; no such sequences were found in the Em2 chromosome and the other 5 *E. meningoseptica* strains (S5 Table). The analyses unveiled that Em1 had a directed repeat (47-bp, GTTGTGCAGTATCACAAATATACTGTAAAATGAAAGCAGTTCACAAC) and had 40 spacers (S5 Table). CRISPR sequences were completely conserved in Em3 (coverage 100%, identity 100%). By scanning the other selected *E. meningoseptica* genomes in the NCBI database, *E. meningoseptica* strain 4076 (CP016376) and NBRC 12535 had a predicted CRISPR (DR length: 47, number of spacers: 21). Em1and Em3 genomes carried only one *cas* gene (type II CRISPR RNA-guided endonuclease Cas9) flanking the CRISPR regions. The amino acid sequences of Cas9 in Em1 showed the highest identity (68%) to that in *Capnocytophaga* spp (data not shown). Only incomplete prophage elements were found in Em1 (2), Em2 (1), and Em3 (2). Moreover, most of these *E. meningoseptica* strains did not have the complete prophages except strain CSID_3000515919 (S6 Table).

## Discussion

*E. meningoseptica* isolated from hospitalized patients in Michigan carried many virulence factors participating in biofilm formation, proteases, lipooligosaccharide (LOS) synthesis, iron uptake and transportation, heat shock proteins, and capsule formation, highlighting that they have great potential to invade and colonize animal hosts. Further, our results also showed that these *E. meningoseptica* strains had several uncommon features including multi-drug resistance, sialic acid synthesis and transportation, and curli formation. Comparative genome analysis also showed that *E. meningoseptica* differed from *E. anophelis* or *E. miricola* in many ways. For example, the average GC content in *E. meningoseptica* was higher than in *E. anophelis* or *E. miricola* (Table 1), highlighting that *E. meningoseptica* evolves differently from *E. anophelis* or *E. miricola*. The predicted functional genes indicated that representative *E. meningoseptica* (Em2) shared less than 86% of the total encoding genes with *E. anophelis* strains and *E. miricola*. In particular, the divergence of *E. meningoseptica* from the other *Elizabethkingia* species was revealed by the different prophages, virulence factors and CRISPR-Cas systems.

By studying how pan-genome size increases as a function of the number of genomes sampled, we can gain insight into a species’ genetic repertoire [25]. Regression analysis of the pan genomes of *E. meningoseptica* showed that the pan-genome of *E. meningoseptica* is evolving through the loss or gain of a range of genes since their divergence as those reported in many pathogens including *E. anophelis*, *Flavobacterium psychrophilum* and *F. spartansii* [16, 34–36]. It is not surprising that *E. meningoseptica* has an open pan-genome because it lives in diverse ecological niches (both aquatic and terrestrial environments), and colonizes and multiplies in animal and plant hosts [1, 37, 38].

*E. meningoseptica* harbored an extensive array of specialized CAZymes (up to 117) for the metabolism of glucans with up to 58 GHs, congruent with metabolic capacity to utilize various carbon sources. Some of the GH genes are organized like the typical PULs that are widely found in Bacteroidetes [39–41]. Differing from aquatic bacteria, terrestrial flavobacteria always carry diverse CAZymes with a large potential for the breakdown of polysaccharides from plants or animals [41–43]. Instead, less CAZyme genes are expected in the genomes of marine flavobacteria because they usually utilize peptides (rhodopsins) and/or harvesting light under the nutrient stressing environment [41, 44, 45]. The CAZyme components and assembly patterns are important properties that reveal *Elizabethkingia* may utilize a variety of carbon sources [4, 46–49]. These results further support the notion that *Elizabethkingia* can adapt to environmental variations between the terrestrial and aquatic native habitats they colonize [16, 18, 46]. Besides soil and water, *Elizabethkingia* are often associated with the amphibious animals such as frogs and insects (i.e. mosquitoes) living in both aquatic and terrestrial environments [49, 50].

The detailed mechanisms for *Elizabethkingia* transmission from the environment to healthcare facilities and to patients remain unclear [18]. However, infection by *Elizabethkingia* can be mediated by multiple pathways of exposure [18]. Regardless of the transmission pathways, the genes in *E. meningoseptica* strains involved in biofilm on the abiotic material are one of the most important virulence factors [1]. The biofilm allows bacteria to persist on the surface of the medical utilities and resist against disinfection reagents [1]. Moreover, attachment/adhesion to the external surfaces and/or tissues of animal hosts is critical in the course of *E. meningoseptica* infection [1, 36, 51]. Many virulence factor genes including liposaccharide, hemagglutinin, capsule, and curli formation are conserved in *E. meningoseptica* (S1 Table). Capsule components are well known to play a vital role in the adhesion and/or biofilm formation [16]. A previous study has shown that expression of hemagglutinin adhesins allowed bacterial attachment and subsequent cell accumulation on target substrates [52]. Curli fibers not only participate in biofilm formation but also contribute to bacterial adhesion to animal cells [52]. *E. meningoseptica* Em1, Em2 and Em3 had hydrophilic cell surfaces. However, it remains unclear if such features affect bacterial adhesion ability [1]. Jacobs and Chenia (2011) investigated biofilm formation and adherence characteristics of *E. meningoseptica* isolated from freshwater tilapia [1]. The results showed that *E. meningoseptica* displayed better biofilm formation in nutrient-rich medium than that in nutrient-limited one at both low and high temperature [1].

Sialic acid is generally found at the terminal position(s) within glycan molecules covering animal cell surfaces [53]. Sialic acid is involved in various cellular processes including intercellular adhesion, cell signaling and immune system invasion [54]. In bacteria, sialic acid molecules participate in many pathogenesis processes [54–56]. For instance, it can be integrated into cell components (i.e. the membrane lipopolysaccharide and capsule) that mimic the surface molecules of the host cell (called molecular mimicry), and thereby assist pathogens to escape the host innate immune response [56]. Moreover, sialic acid is also a good carbon as well as nitrogen source when environmental nutrients are limited [57]. Sialic acid enters bacteria through transporters and is next converted into fructose-6-phosphate; thus, it enters the central metabolism pathway [57, 58]. Additionally, sialic acid-rich glycolipids, glycoproteins and proteoglycans maintain water molecules at the bacterial surface, contributing to the uptake of polar molecules [56]. The detailed physiological roles of sialic acid remain unknown in flavobacteria. However, some *E. meningoseptica* strains (G4120, PRCM and CSID_3000515919) potentially acquire sialic acid by either uptake or the *de novo* synthesis pathway (Table 5). Other *Elizabethkingia* may only utilize sialic acid by *de novo* synthesis because they lack the exosialidase and sialic acid uptake machinery (Table 5). *G. vaginalis* carrying the putative sialidase A gene (*nanH*) was associated with the presence of vaginal biofilms, indicating that sialidase A is an important virulence factor for bacterial vaginosis (BV) [59]. *nanH* mutant failed to form biofilms in *T. forsythia* [60]. However, it restored the biofilm when purified NanH protein was added, showing that sialidase A was critical for pathogens to acquire nutrient source [60]. The same study further suggested that sialidase inhibitors might be useful adjuncts in periodontal therapy. In this study, genes encoding sialidase A (*nanH*) or sialic acid transporter genes (*neu*C1 and C2) were absent from *E. miricola* (except strain CSID_3000517120) and *E. anophelis*, indicating that different *Elizabethkingia* may display different pathogenesis mechanisms.

Antimicrobial susceptibility pattern for *Elizabethkingia* is often controversial [3, 18, 61]. Therefore, the selection of appropriate empiric therapy early in the course of bacterial infection can be very challenging. For example, Han et al [13] reported that 100% of *E. meningoseptica* isolates from South Korea were susceptible to piperacillin-tazobactam and less susceptible to ciprofloxacin (23% of isolates). However, *E. meningoseptica* clinical isolates from Taiwan were resistant to piperacillin/tazobactam while 74.4% of the strains were susceptible to trimethoprim/sulfamethoxaz [7]. The antibiotic susceptibility patterns in Michigan isolates mimic those isolates from Taiwan. Such discrepancies indicate that *E. meningoseptica* from different geographical regions may evolve different antibiotic resistance mechanisms [62]. Thus, the investigation of the molecular mechanisms involving in antibiotic-resistance is required, which could directly contribute to the empirical treatment of *E. meningoseptica* infections [62, 63]. Remarkably, the genomes of *E. meningoseptica* carried intrinsic class A extended-spectrum β-lactamases (ESBLs) and inherent class B metallo-β-lactamases (MBLs) [15]. These genes are likely to confer resistance to these β-lactam antibiotics including the combination drug piperacillin/tazobactam (Table 3). *E. meningoseptica* genomes shared many antibiotic-resistance genes with those from other *Elizabethkingia* species while there was minor differences [14, 64]. Besides those antibiotic-inactivating enzymes/proteins, *Elizabethkingia* may utilize the multidrug efflux pumps to excrete a wide range of antibiotics [65], including *abes S*, *msrB*, *macB*, *ceoB*, *adeG* and many others (Table 4). For example, CeoB, a component of RND multidrug efflux system, pumps out chloramphenicol and ciprofloxacin [65, 66]. MsrB, as an ABC-efflux pump, was reported in *Staphylococcus* species to confer resistance to erythromycin and streptogramin B antibiotics [67]. Genes involving in tetracycline and vancomycin resistance were detected in *Elizabethkingia* in our study and others [64], which may explain why these drugs are not very effective in eliminating *Elizabethkingia* infections. Analysis of the antibiotic resistance in *E. meningoseptica* showed that they carried genes conferring to drug resistance including aminoglycosides, isoniazid, streptogramin, aminosalicylate, fluoroquinolone, macrolide, tetracycline, diaminopyrimidine, phenico, glycopeptide and rifamycin. Instead, we did not detect genes involving in resistance of sulfonamide (folate pathway inhibitors) in the genomes of Michigan isolates, which agrees with their susceptibility to trimethoprim/sulfamethoxazole. Mutation of the *gyrB* gene encoding the DNA gyrase subunit B in strains Em1, Em2 and Em3 was predicted to contribute to fluoroquinolones’ resistance (such as ciprofloxacin) in this study.

Prophages are known to modulate virulence and antibiotic resistance gene expression, which alters the production and/or secretion of toxins during the infection course [68, 69]. Further, they are one of the important contributors to genetic diversification by transferring the functional genes among the different strains [70]. The rare occurrence of complete prophages in *E. meningoseptica* remains unclear. Our analysis revealed only few of the selected genomes (Em1 and Em3) had “confirmed” CRISPRs (Supplementary Table S6). The similar observations were reported in *E. anophelis* genomes [16]. The defense of the invasions of foreign genetic elements such as plasmids, transposons or phages may require both restriction modification systems (RMs) and Clustered Regularly Interspaced Short Palindromic repeat sequences (CRISPRs) in *Elizabethkingia* [71]. However, the detailed mechanisms need to be further investigated.

## Acknowledgments

This project was funded by NIH grant R37AI21884.

## Contributions

**Conceptualization:** Shicheng Chen, Marty Soehnlen and Edward D. Walker.

**Data curation:** Shicheng Chen.

**Formal analysis:** Shicheng Chen.

**Funding acquisition:** Edward D. Walker.

**Investigation:** Shicheng Chen, Jochen Blom, Nicolas Terrapon, Bernard Henrissat.

**Methodology:** Shicheng Chen, Jochen Blom, Nicolas Terrapon, Bernard Henrissat.

**Software:** Jochen Blom, Nicolas Terrapon, Bernard Henrissat.

**Supervision:** Shicheng Chen, Marty Soehnlen and Edward D. Walker.

**Validation:** Shicheng Chen.

**Visualization:** Jochen Blom, Nicolas Terrapon, Bernard Henrissat.

**Writing – original draft:** Shicheng Chen, Jochen Blom, Nicolas Terrapon, Bernard Henrissat.

**Writing – review & editing:** Shicheng Chen, Jochen Blom, Nicolas Terrapon, Bernard Henrissat.

