## Supplemental Figures for "Comparative genomic analyses reveal diverse virulence factors and antimicrobial resistance mechanisms in clinical *Elizabethkingia meningoseptica* strains"

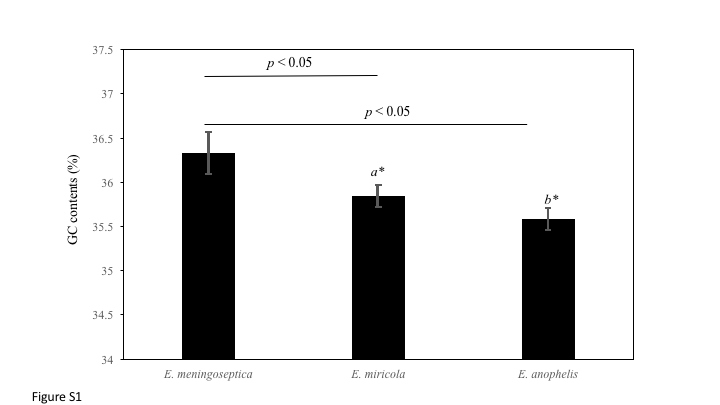

**Figure S1. Comparisions of the GC contents in *E. meningoseptica, E. anophelis* and *E. miricola***. The selected *E. meningoseptica* strains are those listed on the NCBI database (cutoff date 11/15/2018): <https://www.ncbi.nlm.nih.gov/genome/?term=Elizabethkingia>.

**A**

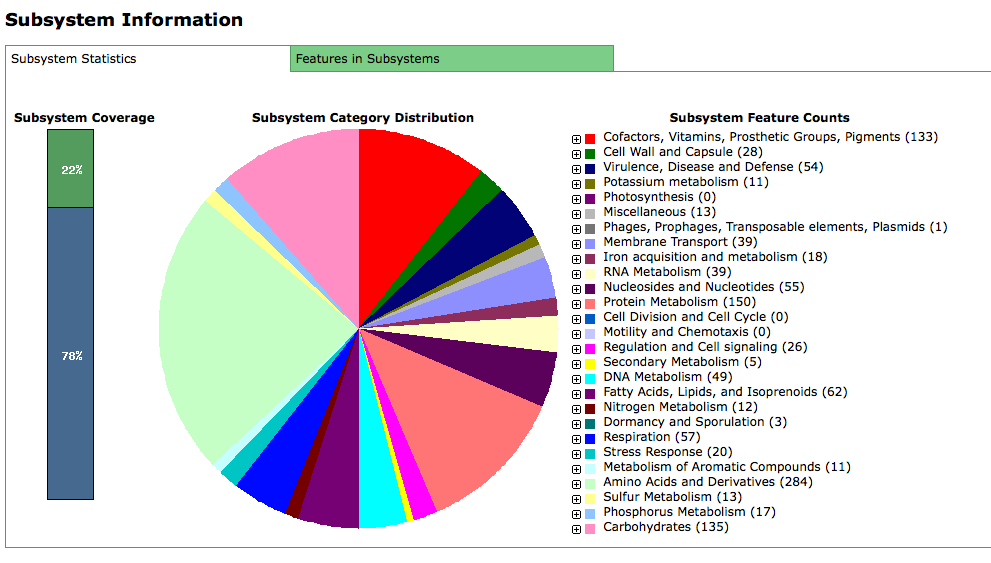

**B**

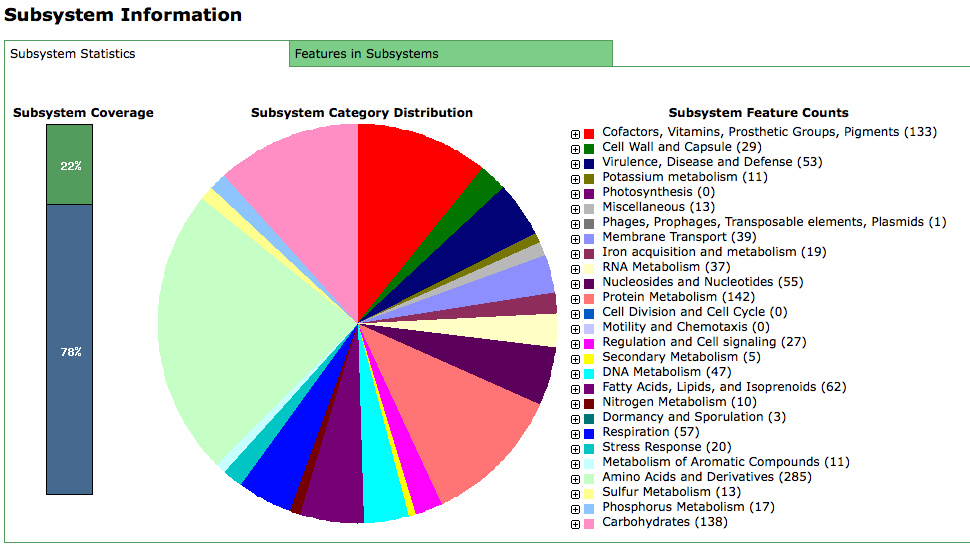

**C**

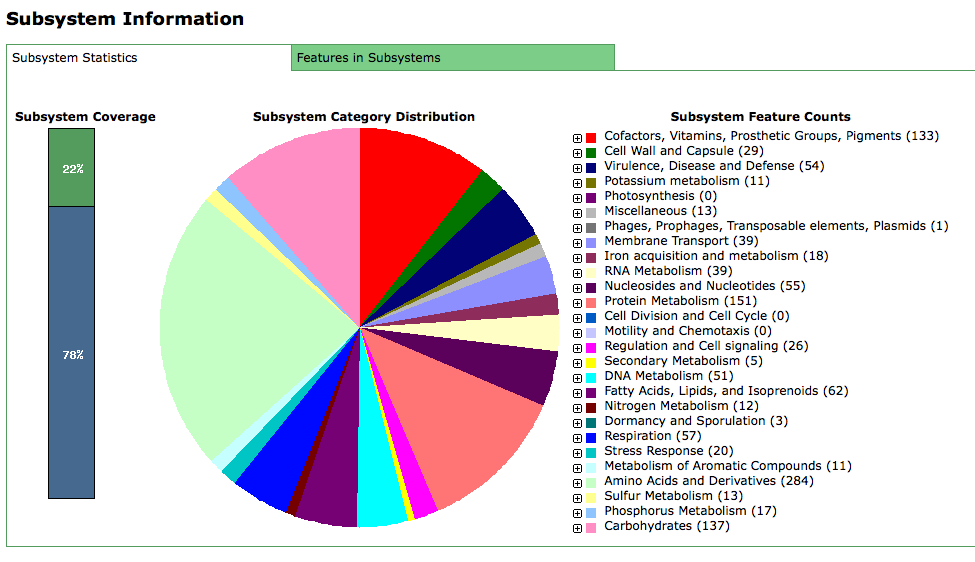

**Figure S2. RAST analysis of the selected *E. meningoseptica* Em1 (A), Em2 (B) and Em3 (C).**

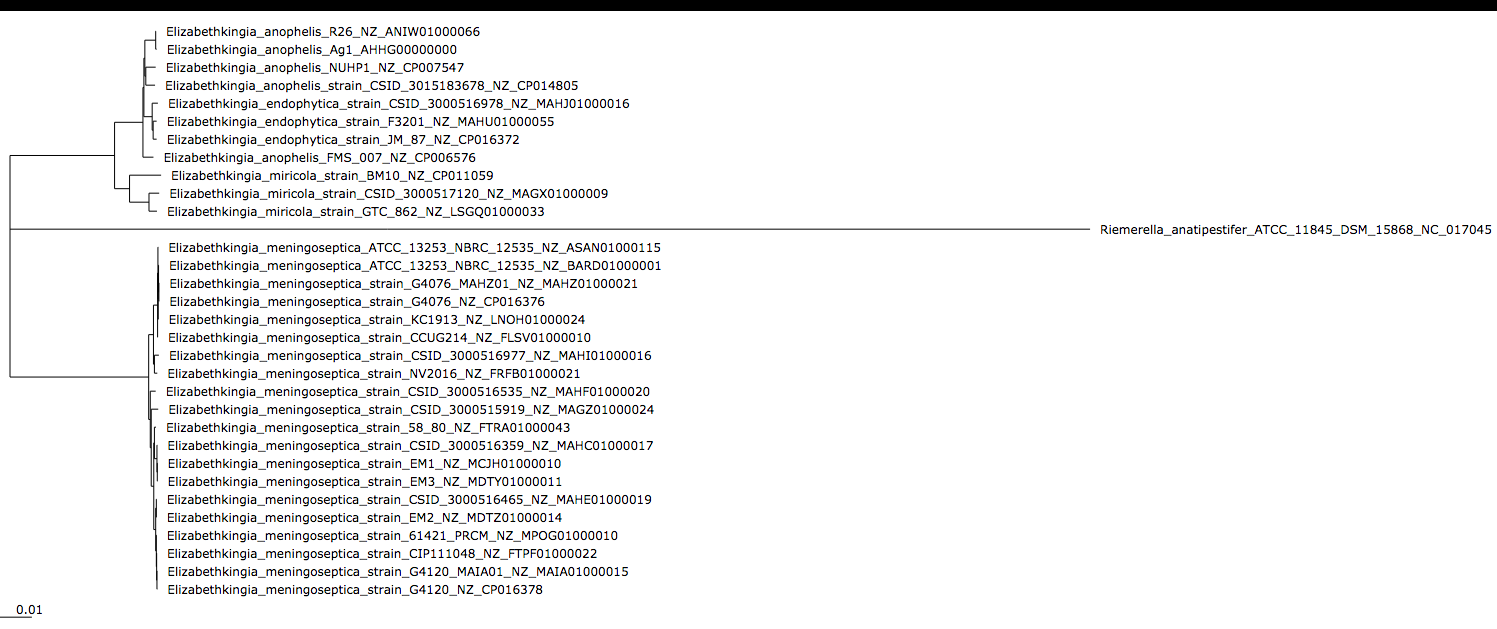
**Figure S3. Phylogenetic relationship among the selected *Elizabethkingia*.** Tree was constructed for 32 genomes with a core of 1170 genes per genome, 37440 in total. The core has 405494 AA-residues/ bp per genome, 12975808 in total.

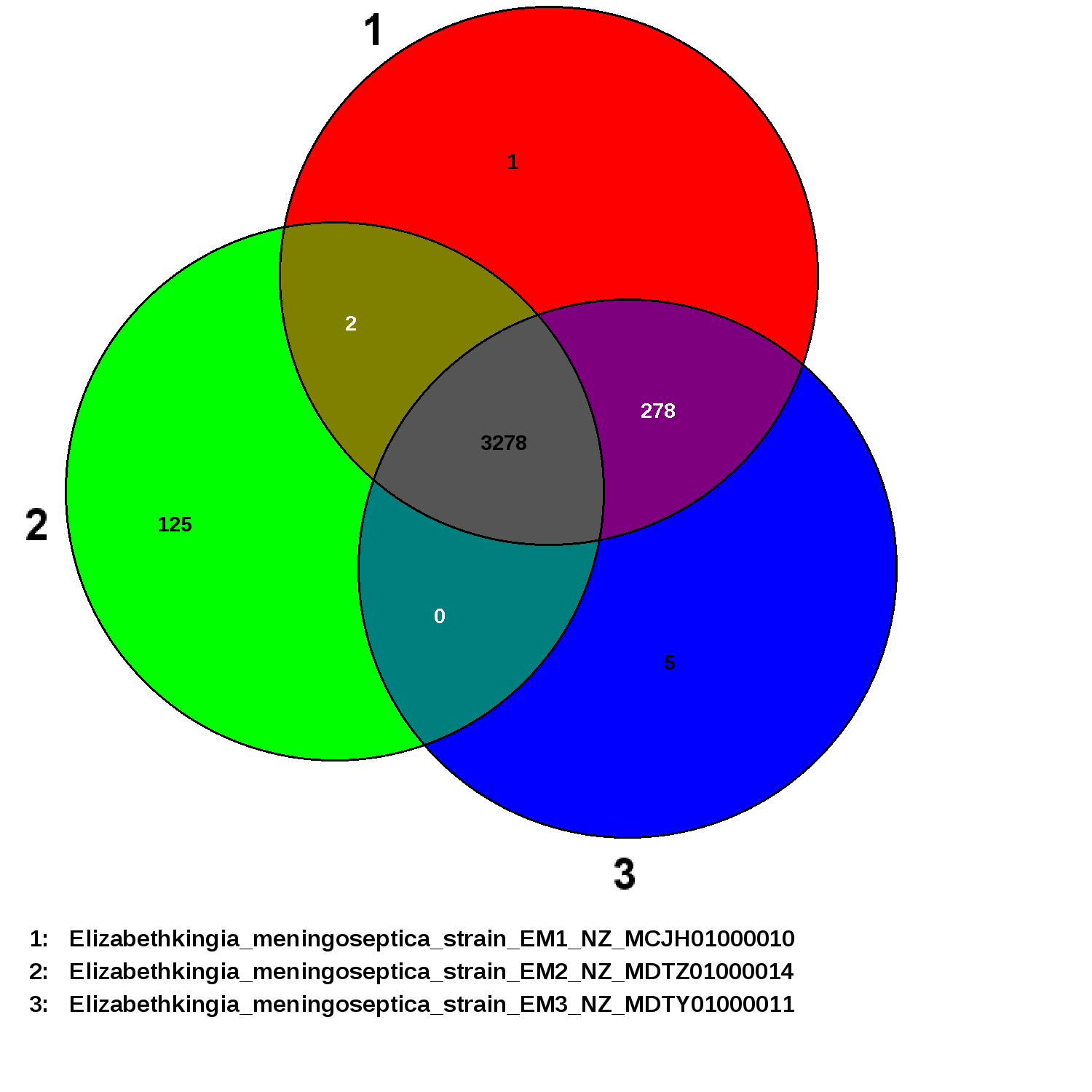

**Figure S4. The shared genes among the selected *E. meningoseptica* Em1, Em2 and Em3.** EDGAR was used for Venn diagrams.

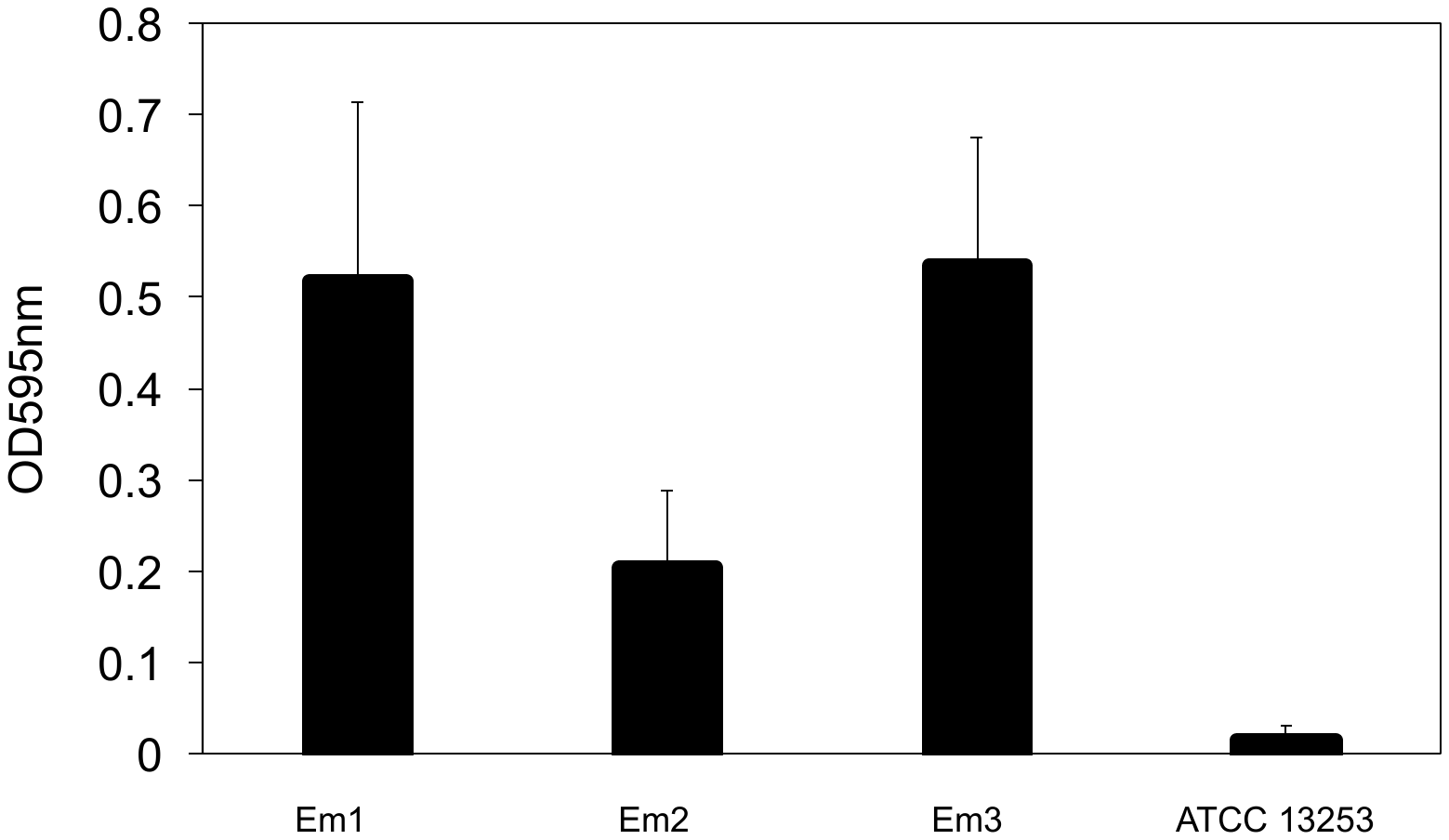

**Figure S5. *In vitro* biofilm assay in the selected *E. meningoseptica*.** The cells were cultured by shaking in TSB at 37 °C to obtained the initial inocula and the cell density was adjusted to the same OD at 600 nm (0.1). 200 μl were inoculated on 96-well plates for at least 24 hours. The biofilm assay was carried out using crystal blue staining.
