## Supplementary material for "Comparative genomic analyses reveal diverse virulence factors and antimicrobial resistance mechanisms in clinical *Elizabethkingia meningoseptica* strains": S1 Table

**Table S1. Comparations of the predicted virulence factors among various *Elizabethkingia***

|  | *E. meningoseptica* | | | | | | | | | *E. anophelis* | | | *E. miricola* | |
| --- | --- | --- | --- | --- | --- | --- | --- | --- | --- | --- | --- | --- | --- | --- |
| Gene products | Em2 | Em1 | Em3 | 5880 | CSID 3000516977 | G4120 | 61421 PRCM | NBRC 12535 | Ag1 | | NHU1 | BM10 | | CSID_3000517120 |
| DnaJ | 100 | 100 | 100 | 100 | 100 | 100 | 100 | 100 | 91 | | 91 | 91 | | 90 |
| 30S ribosomal protein S9 | 100 | 100 | 100 | 100 | 100 | 100 | 100 | 100 | 99 | | 99 | 97 | | 97 |
| 50S ribosomal protein L13 | 100 | 100 | 100 | 100 | 100 | 100 | 100 | 100 | 97 | | 97 | 96 | | 96 |
| ABC transporter ATP-binding protein | 100 | 100 | 100 | 100 | 100 | 100 | 100 | 100 | 96 | | 96 | 96 | | 96 |
| Alcohol dehydrogenase | 100 | 100 | 100 | 100 | 99 | 100 | 99 | 99 | 96 | | 96 | 96 | | 96 |
| Aldehyde dehydrogenase | 100 | 100 | 99 | 100 | 99 | 100 | 100 | 100 | 95 | | 95 | 96 | | 96 |
| Aldehyde reductase | 100 | 100 | 99 | 100 | 98 | 99 | 99 | 99 | 92 | | 93 | 93 | | 93 |
| Alpha-mannosidase | 100 | 99 | 99 | 99 | 99 | 99 | 99 | 99 | 91 | | 91 | 91 | | - |
| Alpha-mannosidase 2 | 100 | 100 | 99 | 100 | 100 | 100 | 100 | 100 | 94 | | 94 | 94 | | 94 |
| Asparagine--tRNA ligase | 100 | 100 | 100 | 100 | 100 | 100 | 100 | 100 | 95 | | 95 | 95 | | 95 |
| Aspartate 1-decarboxylase | 100 | 100 | 100 | 100 | 100 | 100 | 100 | 100 | 98 | | 98 | 99 | | 99 |
| Bla | 100 | 98 | 98 | 97 | 98 | 100 | 100 | 100 | 77 | | 75 | 75 | | 75 |
| Capsule biosynthesis protein CapD | 100 | - | - | - | - | 100 | 100 | 100 | - | | - | - | | 61 |
| Catalase | 100 | 99 | 99 | 99 | 99 | 99 | 99 | 99 | 93 | | 93 | 93 | | 94 |
| Catalase/peroxidase | 100 | 100 | 99 | 100 | 99 | 100 | 100 | 100 | 90 | | 90 | 89 | | 89 |
| Chloramphenicol O-acetyltransferase | 100 | 98 | 97 | 98 | 97 | 100 | 100 | 100 | 86 | | 86 | 87 | | 87 |
| ClbS/DfsB | 100 | 100 | 95 | 100 | 95 | 100 | 100 | 100 | 74 | | 74 | 73 | | 75 |
| Clp2 | 100 | 100 | 100 | 100 | 100 | 100 | 100 | 100 | 96 | | 96 | 96 | | 96 |
| ClpX | 100 | 100 | 99 | 100 | 100 | 100 | 100 | 100 | 95 | | 95 | 95 | | 95 |
| CTP synthetase | 100 | 100 | 100 | 100 | 100 | 100 | 100 | 100 | 99 | | 98 | 98 | | 98 |
| Cysteine sulfinate desulfinase | 100 | 99 | 99 | 99 | 99 | 100 | 100 | 100 | 93 | | 93 | 93 | | 93 |
| DNA starvation/stationary phase protection protein | 100 | 100 | 100 | 100 | 100 | 100 | 100 | 100 | 97 | | 96 | 96 | | 96 |
| DNA-binding protein | 100 | 100 | 100 | 100 | 100 | 100 | 100 | 100 | 100 | | 100 | 98 | | 98 |
| DNA-directed RNA polymerase subunit beta | 100 | 100 | 100 | 100 | 100 | 100 | 100 | 100 | 98 | | 98 | 98 | | 98 |
| DNA-directed RNA polymerase subunit beta | 100 | 100 | 100 | 100 | 100 | 100 | 100 | 100 | 99 | | 99 | 99 | | 99 |
| DnaK | 100 | 100 | 100 | 100 | 99 | 99 | 99 | 99 | 97 | | 97 | 96 | | 96 |
| dTDP-glucose 4,6-dehydratase | 100 | 98 | 98 | 100 | 98 | 100 | 100 | 100 | 95 | | 96 | 94 | | 94 |
| Glucose-1-phosphate thymidylyltransferase | 100 | 99 | 96 | 100 | 96 | 100 | 100 | 100 | 95 | | 94 | 94 | | 94 |
| Glutamate dehydrogenase | 100 | 100 | 100 | 100 | 99 | 100 | 100 | 100 | 98 | | 98 | 98 | | 98 |
| GroL | 100 | 100 | 99 | 100 | 99 | 100 | 100 | 100 | 99 | | 98 | 98 | | 98 |
| HxlR | 100 | - | - | - | - | 100 | 100 | 100 | - | | - | - | | - |
| Hydroxyisourate hydrolase | 100 | 100 | 99 | 100 | 100 | 100 | 100 | 100 | - | | - | - | | - |
| Hypothetical protein | 100 | 99 | 99 | 99 | 99 | 99 | 99 | 99 | 69 | | 70 | 68 | | 70 |
| Hypothetical protein 2 | 100 | 100 | 98 | 100 | 98 | 100 | 100 | 100 | 60 | | 90 | 91 | | 90 |
| Hypothetical protein 3 | 100 | 100 | 99 | 100 | 99 | 100 | 100 | 100 | 86 | | 85 | 85 | | 83 |
| Isocitrate lyase | 100 | 98 | 100 | 98 | 100 | 100 | 100 | 100 | 92 | | 92 | 92 | | 92 |
| Methylglyoxal synthase | 100 | 100 | 99 | 99 | 99 | 100 | 100 | 100 | 97 | | 96 | 94 | | 94 |
| Pantetheine-phosphate adenylyltransferase | 100 | 100 | 100 | 100 | 100 | 100 | 100 | 100 | 94 | | 94 | 94 | | 94 |
| Peptidylprolyl isomerase | 100 | 100 | 100 | 100 | 100 | 100 | 100 | 100 | 95 | | 95 | 93 | | 95 |
| Phosphomethylpyrimidine synthase ThiC | 100 | - | - | - | - | 100 | 100 | 100 | - | | - | - | | - |
| Short-chain dehydrogenase | 100 | 99 | 98 | 100 | 98 | 100 | 100 | 100 | 89 | | 88 | 88 | | 89 |
| Sigma-54-dependent Fis family transcriptional regulator | 100 | 100 | 100 | 100 | 100 | 100 | 100 | 100 | 93 | | 93 | 94 | | 94 |
| SOD1 | 100 | 99 | 99 | 99 | 99 | 98 | 98 | 98 | 84 | | 84 | 84 | | 84 |
| SOD2 | 100 | 100 | 99 | 100 | 99 | 100 | 100 | 100 | 96 | | 96 | 96 | | 96 |
| SOD3 | 100 | 100 | 100 | 100 | 100 | 100 | 100 | 100 | 88 | | 88 | 88 | | 88 |
| TF-1 | 100 | 100 | 100 | 100 | 100 | 100 | 100 | 100 | 98 | | 98 | 98 | | 98 |
| Thioredoxin-disulfide reductase | 100 | 100 | 100 | 100 | 99 | 100 | 100 | 100 | 96 | | 96 | 95 | | 95 |
| Transporter | 100 | 99 | 100 | 99 | 98 | 100 | 100 | 100 | 59 | | 54 | 59 | | 59 |
| UDP-glucose 6-dehydrogenase | 100 | - | 99 | 99 | 99 | 100 | 100 | 100 | 87 | | 87 | 86 | | 86 |
