## Supplementary material for "Comparative genomic analyses reveal diverse virulence factors and antimicrobial resistance mechanisms in clinical *Elizabethkingia meningoseptica* strains": S2 Table

|  |  | **Table S2 Comparative analysis of the putative biofilm formation in *Elizabethkingia* spp.** | | | | | | | | | | | | | | |
| --- | --- | --- | --- | --- | --- | --- | --- | --- | --- | --- | --- | --- | --- | --- | --- | --- |
|  |  |  | ***E. meningoseptica*** | | | | | | | | ***E. anophelis*** | | | | ***E. miricola*** | |
|  | **Accession number** | **Genes** | **Em2** | **Em1** | **EM3** | **5880** | **CSID**  **3000516977** | **G4120** | **61421 PRCM** | **NBRC 12535** | **Ag1** | **NUHP1** | **CSID**  **3015183678** | **BM10** | | **CSID**  **3000517120** |
| Capsule formation |  |  |  |  |  |  |  |  |  |  |  |  |  |  | |  |
|  |  | *cap8E* | 100 | - | - | - | - | 100 | 100 | - | - | - | - | - | | 96 |
|  | WP_070904261 | *cap8G* | 100 | - | - | - | - | 100 | 100 | - | - | - | - | - | | 96 |
|  | WP_070904247 | *cap8O* | 100 | - | - | - | - | 100 | 100 | - | - | - | - | - | | - |
|  | WP_070904244 | *cap8D* | 100 | 99 | 99 | - | 99 | 100 | 100 | 99 | 95 | 95 | 95 | 95 | | 94 |
|  | WP_070904262 | *cap4F* | 100 | - | - | - | - | 100 | 100 | - | - | - | - | - | | 90 |
|  | WP_070904260 | *cap8F* | 100 | - | - | - | - | 100 | 100 | - | - | - | - | - | | 88 |
|  | WP_070904246 | *cps4D* | 100 | 82 | 82 | - | 82 | 100 | 100 | 82 | 81 | 83 | 83 | 81 | | 79 |
|  | WP_070904232 | *Cj1137c* | 100 | 99 | 99 | - | - | 100 | 100 | - | - | - | 99 | - | | 69 |
|  | WP_070904673 | *capD* | 100 | 99 | 99 | 99 | 99 | 100 | 100 | 99 | 93 | 93 | 93 | 94 | | 93 |
|  | WP_070904243 | *cap4D* | 100 | 96 | 96 | - | 83 | 100 | 100 | 84 | 79 | 76 | 77 | 73 | | 77 |
| Curli | WP_069214179 | *CurEm1* | 100 | 100 | 100 | 100 | 99 | 100 | 100 | 98 | - | - | - | - | | - |
|  | WP_069214180 | *CurEm2* | 100 | 100 | 100 | 100 | 99 | 100 | 100 | 99 | - | - | - | - | | - |
|  | WP_069214181 | *CurEm3* | 100 | 100 | 100 | 100 | 99 | 100 | 100 | 99 | - | - | - | - | | - |
|  | WP_070904486 | *CurEm4* | 100 | 99 | 99 | 99 | 99 | 100 | 100 | 99 | - | - | - | - | | - |
| Flagellar motor |  |  |  |  |  |  |  |  |  |  |  |  |  |  | |  |
|  | WP_069215025 | *FlgD* | 100 | 100 | 100 | 100 | 100 | 100 | 100 | 99 | 89 | 89 | 100 | 90 | | 89 |
|  | WP_070904468 | *MotB* | 100 | 99 | 99 | 99 | 99 | 99 | 99 | 99 | 87 | 87 | 87 | 87 | | 83 |
